# Rapid Semantic Processing: An MEG Study of Narrative Text Reading

**DOI:** 10.1101/2024.11.27.625607

**Authors:** Anastasia Neklyudova, Gurgen Soghoyan, Olga Martynova, Olga Sysoeva

## Abstract

Research has shown that semantic analysis occurs at early stages of word processing (less than 200 ms). While traditional studies have focused on isolated words/sentences, our research explores rapid semantic processing during reading stories using event-related potentials and magnetoencephalography. We employed the rapid serial visual presentation paradigm to present texts word by word. Each word presentation lasted 150 ms, enhancing rapid semantic processing. We computed semantic dissimilarity (SD) values for each word and categorized them into quartiles to investigate their effect on brain responses. Our analysis revealed significant ERP differences within early time windows (120-132 ms). Two distinct clusters were identified: positive in the right occipital region and negative in the left temporal region. In both clusters less pronounced responses were registered for words with lowest SD which corresponds to theories of predictive coding. These findings broaden our understanding of rapid semantic processing and suggest new methodology.

## Introduction

Reading is a unique human ability that enables us to use visual symbols to understand the meaning of words. Children learn to read at 6-8 years, and the speed of their reading increases drastically when they achieve the level of proficiency (Korinth & Nagler, 2021). For a skilled adult reader, the brain can extract information from a written word in less than half a second. Eye-tracking studies have shown that fixation time on each word is approximately 200 ms and remains consistent across different languages (Gagl et al., 2022).

A major debate in cognitive science concerns whether different levels of language information are processed serially or in parallel. Traditionally, reading involves distinct stages: first, visual-orthographic processing and mapping orthography to phonology, then lexical-semantic processing (Bentin et al., 1985; Holcomb & Grainger, 2007; Pulvermüller et al., 2009). In contrast, parallel models suggest that these types of information are accessed simultaneously since the early stage of processing (Pulvermüller et al., 2009).

Event-related potentials (ERPs) are a powerful method to show how brain activity supporting reading develops in time. The N400 and P600 are probably the most extensively studied ERP components in language research. The N400, which peaks 300-500 ms after a word’ onset, was first associated with semantically incongruent words (Kutas & Hillyard, 1980) and has been found to be larger for meaningless pseudowords and rare words (Kutas & Federmeier, 2000; Van Petten & Kutas, 1990). The P600 is typically associated with the processing of both semantic and syntactic anomalies or incongruities in the language (Gouvea et al., 2010; Osterhout & Holcomb, 1992; van Herten et al., 2005).

ERP studies show that, within 100-250 ms, brain responses to visual words are influenced by word frequency, length, semantics, and emotional properties (Amsel et al., 2013; Assadollahi & Pulvermüller, 2001; Hauk et al., 2006; Kim & Lai, 2012; Scott et al., 2009; Skrandies, 1998). Other research has shown even earlier responses to semantic features, for example the distinction between abstract and concrete words occurring at 40-100 ms (Sysoeva et al., 2007) and the differentiation between words and pseudowords at 50-80 ms (MacGregor et al., 2012; Shtyrov & MacGregor, 2016).

The most common approach to studying speech processing is to present individual words or sentences with various semantic or lexical features, often using the rapid serial visual presentation (RSVP) paradigm to show sentences word-by-word. This approach helps avoid contamination of ERPs by eye movements, which is crucial for studying early components, while also providing a reading experience that is closer to natural reading situations. Although several aspects of word processing differ between RSVP and natural reading (Kornrumpf et al., 2016), all ERP components are preserved and have been reported to be modulated by semantic and lexical features.

However, there is a growing trend nowadays in using natural stimuli (i.e., complete narrative stories) to study spoken and written speech processing (Hamilton & Huth, 2020). Typically, brain responses to these stimuli are analyzed using neural tracking tools (e.g., coherence, temporal response functions). Here, we introduce the use of event-related potentials (ERP) to analyze MEG responses to narrative stories. We believe the ERP technique is especially valuable for studying rapid brain responses, as it directly reflects the temporal evolution of information processing.

This study used the RSVP paradigm to examine rapid semantic processing during reading narrative stories. We recorded MEG data, which offers EEG’s high temporal resolution with better localization of brain responses. By presenting words for only 150 ms, we aimed to explore the brain ability for quick language analysis. Instead of selecting specific words for semantic processing, we computed semantic dissimilarity (SD) as distributed values for all presented words and then averaged the words with low and high SD for ERP analysis. We hypothesize that semantic dissimilarity will be reflected in brain activity already within the first 150 ms after stimuli presentation.

## Materials and Methods

### Procedure

Twenty-seven adults (17 females) participated, with an average age of 26.32 ± 4.48 years (range: 18.96-40.43). All were right-handed, native Russian speakers with normal or corrected vision, and no reported neurological or psychiatric issues. Written consent was obtained after explaining the procedures. The study, approved by the Institute of Higher Nervous Activity and Neurophysiology’s ethics committee (June 30, 2023), followed the Helsinki Declaration’s ethical guidelines for human research.

Participants sat in a comfortable chair in a magnetically shielded room, reading text projected by a PT-D7700E-K DLP projector (1280×1024 resolution, 60 Hz). Six stories in Russian, totaling 2,533 words, were presented word-by-word with Psychopy software, each word shown for 150 ms without pauses. Words combined with punctuation (e.g., periods, commas) were excluded from analysis. The reading session, with breaks, lasted about 20 minutes, while the full recording session, including other tasks and preparation, took approximately 2.5 hours. During breaks, participants answered 14 open-ended comprehension questions. Responses were rated by experimenters on a 3-point scale: 2 for correct, 1 for minor errors, and 0 for incorrect or no response, with a maximum possible score of 20.

### MEG processing

MEG data were collected at the Moscow Center for Neurocognitive Research using an Elekta VectorView 306-channel MEG system (Helsinki, Finland) with built-in filters of 0.03–330 Hz and a sampling frequency of 1000 Hz. The system included 102 magnetometers and 204 orthogonal planar gradiometers at 102 positions. Bad channels were identified visually, and the signal was preprocessed with MaxFilter software (version 2.2.15) to reduce external noise using the temporal signal-space separation method (tSSS) and to realign data from different blocks to a standard head position.

Further preprocessing was carried out using MNE-Python software (v.0.24.1). Initially, data from the different recorded stories were combined. Ocular and cardiac artifacts were then removed using Independent Component Analysis (ICA) on the combined data, average number of rejected ICA components was 0.96, range: 0-2.

The artifact-free data were subsequently segmented into epochs from -0.05 to 0.17 seconds relative to each stimulus (i.e., word presentation). For further analysis we used only magnetometers (n=102). Baseline correction was applied from -0.05 to 0 seconds. We excluded epochs where signal amplitude was above 4e-12 T for magnetometers and additionally excluded epochs where amplitude exceeded 5 standard deviations (STD). In two participants the number of deleted epochs varied between 50-93, so we did not include their data to the following analysis. The average number of deleted epochs in the other participants was 1.17±0.3.

### Stimuli preparation

Semantic dissimilarity (SD) was calculated using a word vectorization method called word2vec (Mikolov et al., 2013). This approach calculates semantic dissimilarity (SD) by correlating the vector of the current word with the semantic context of previous words and subtracting this value from 1. The semantic context is estimated as the average of the semantic vectors for all words in the current sentence. For the first words in a sentence, the semantic context is derived from the average of the semantic vectors in the preceding sentence.

We generated a distribution of all SD values, dividing them into quartiles (0.25, 0.5, 0.75, 1.0). Words in the first quartile were labeled ‘low 2,’ in the second ‘low 1,’ in the third ‘high 1,’ and in the fourth ‘high 2.’ These labels were assigned to the corresponding events, and we created epochs based on these four groups for comparison in subsequent analyses.

**Figure 1.**
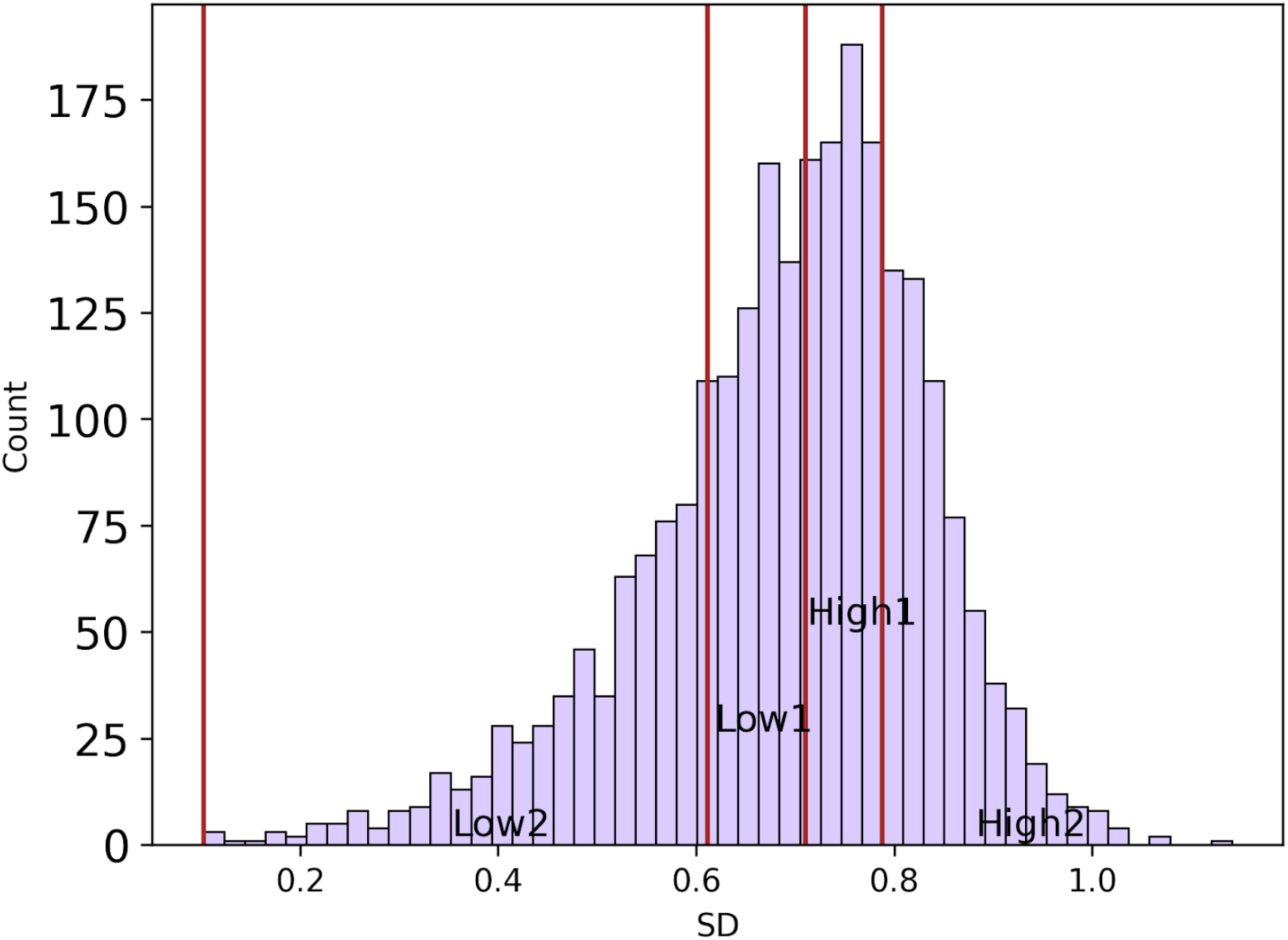
The distribution of semantic dissimilarity (SD) values is shown, with groups of words categorized into four categories based on this metric: low 2 SD, low 1 SD, high 1 SD and high 2 SD.

### Statistical analysis

To identify source clusters of significant differences between conditions (low 2, low 1, high 1, high 2 SD), we employed spatio-temporal cluster analysis. This method addresses multiple comparisons across ERP segments in space and time, allowing for the identification of stable clusters while reducing type I error risk. We set an alpha level of 0.0001 for cluster formation, given the large number of distribution elements. Elements exceeding this threshold were grouped with neighboring elements to form distinct clusters. After conducting 10,000 permutations, we compared the original cluster statistics to the histogram of randomized null statistics using a two-tailed t-test.

## Results

Behavioral assessment indicated an average comprehension questionnaire score of 17.49 (out of 20), with a range of 12-20. All participants reported understanding the content of each story. Individual differences did not correlate with any electrophysiological parameters we studied.

Cluster analysis of ERPs based on semantic dissimilarity (SD) revealed two significantly different sensor clusters. The first cluster, in the right occipital region, showed a significant difference (p=0.001) in the 120-128 ms time window, with lower MEG signal amplitude for the ‘low 2 SD group,’ which differed most from the other three groups. The second cluster, in the left parieto-temporal region, showed a significant difference (p=0.001) in the 125-132 ms window, where the MEG signal amplitude for the ‘low 2 SD’ group was higher than in the other three SD groups.

**Figure 2.**
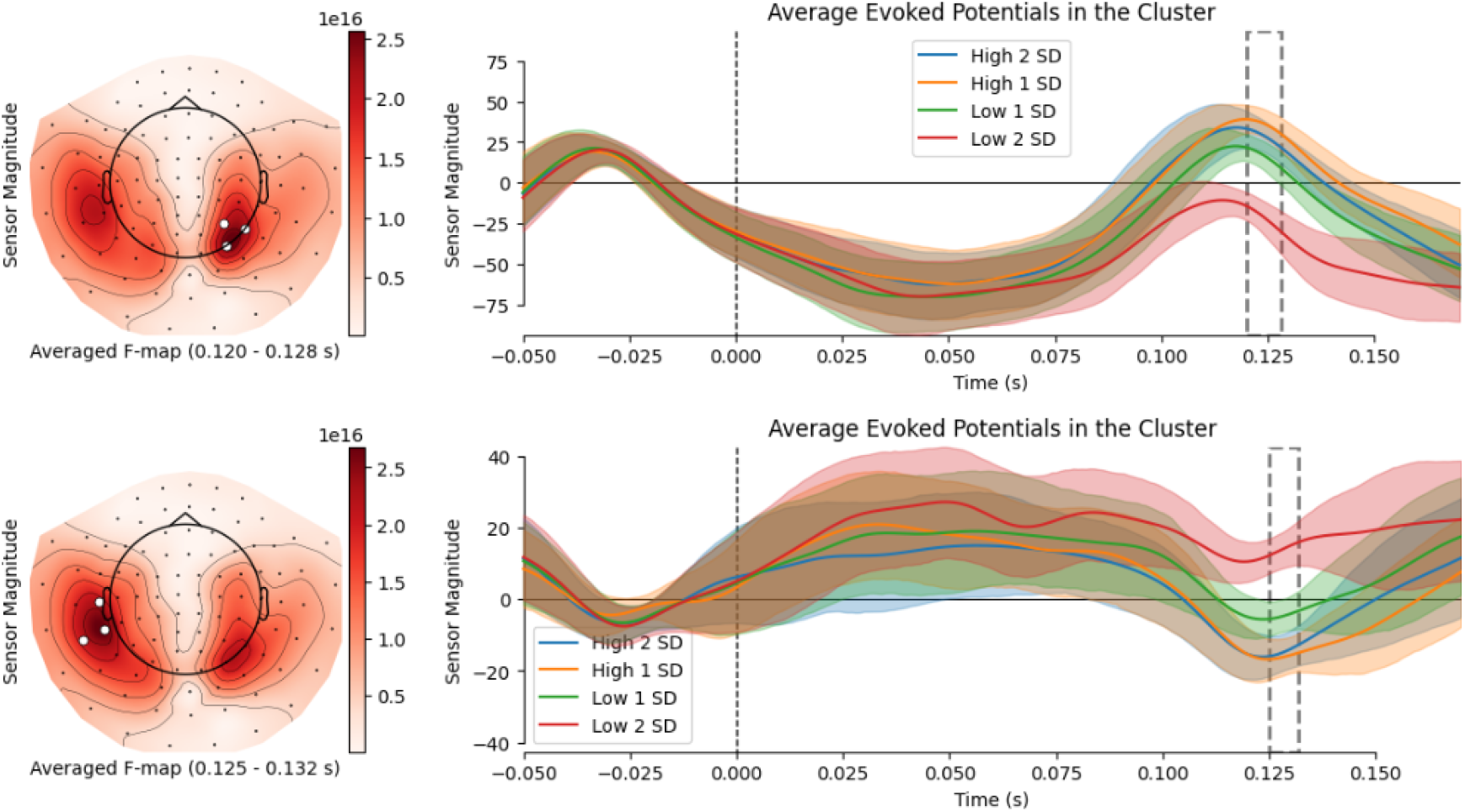
Results of spatio-temporal cluster analysis for words with different rank of semantic dissimilarity (SD): low 2, low 1, high 1 and high 2.

## Discussion

In our study, we analyzed event-related potentials (ERPs) concerning semantic dissimilarity (SD), which enables investigation of semantic processing in context. Previous studies have shown that semantic processing can occur within 200 ms of word presentation (Hauk et al., 2006; MacGregor et al., 2012; Pulvermüller et al., 2009; Sulpizio et al., 2022; Sysoeva et al., 2007). For the first time, we showed rapid semantic processing in brain responses to narrative stories, rather just isolated words or sentences.

Another key aspect of our study was the use of the RSVP paradigm, with a high word presentation rate (150 ms per word), which is faster than the typical 200 ms per word seen in natural reading. This method facilitated rapid semantic processing and showed that comprehension remains unaffected when words are presented individually at this speed.

The difference between words with different rank of SD was evident in the ERPs within early time windows (120-132 ms), revealing two clusters with distinct topographies. The first positive cluster, localized in the right occipital area, showed a weaker response for words with the least SD (low 2 group). The second cluster, with opposite polarity (negative), was found in the left temporal area, where the amplitude for the low 2 group was also lower and most distinct from other SDs. This pattern aligns with the N400 effect, indicating higher amplitudes for semantically unexpected words, consistent with predictive coding theory, which suggests that expected sensory signals elicit attenuated responses (Nour Eddine et al., 2024).

The right medial occipitotemporal gyrus (or the lingual gyrus) has been reported to play a role in rapid automatized naming (RAN), but not in reading, according to an fMRI study by Cummine and colleagues (2015). RAN involves naming highly familiar words as quickly as possible. In our task, each word was presented for only 150 ms. The activation of this region may suggest that rapid semantic processing shares neural networks with RAN.

Studies on dyslexia, characterized by reading difficulties despite preserved intelligence and no sensory issues, often highlight the occipitotemporal, temporoparietal, and left frontal regions as essential for reading (Kronbichler & Kronbichler, 2018). Evidence suggests that the left anterior temporal lobe is involved in semantic processing during both early (∼200 ms) and later (∼400 ms) stages of auditory perception and reading (Bemis & Pylkkänen, 2013; Jackson et al., 2015). Moreover, studies have shown that patients with left temporal lobe epilepsy have altered lexical and semantic processing (Jensen et al., 2011).

This research was funded by the Russian Science Foundation (RSF), grant №23-78-00011. The authors report no competing interests to declare.

